# Behavioral relevance of category selectivity revealed by human ECoG data

**DOI:** 10.1101/2023.12.22.571904

**Authors:** Seyedamin Hashemi, Seyed Ali Ebadi, Fatemeh Zareayan Jahromy, Stefan Treue, Gabriel Kreiman, Moein Esghaei

## Abstract

Object recognition is a crucial brain function that involves a complex interplay between various brain regions. However, the behavioral relevance of functional interactions between these regions remains largely unexplored. In this study, we examined the functional interactions between different brain regions during object recognition using intracranial electrocorticography (ECoG) recordings in subjects diagnosed with pharmacologically intractable epilepsy. We computed the phase locking value (PLV) between different brain areas and its category selectivity, and assessed its behavioral relevance by comparing correctly and incorrectly performed trials. Our results revealed that phase locking between brain regions varies across different object categories and that this variability significantly influences the perceptual behavior of subjects. Importantly, we found that the behavioral relevance of these interactions is spatially organized, with long-range (global) connections being more behaviorally relevant for the frontal lobe and local connections being more crucial for the occipital lobe. These findings underscore the unique roles of different brain areas in object recognition and pave the way for more nuanced explorations of the interplay between brain regions in object recognition and other cognitive functions.

## Introduction

Visual perception of object categories is of crucial importance to the human brain’s function. Many studies have used neural recordings or functional neuroimaging to investigate the underlying neural mechanisms [1]–[5]. Although the ventral visual pathway (especially the inferotemporal (IT) cortex) has been identified to play a key role in identifying objects [1]–[6], there has been considerable debate regarding the extent to which neural information in the temporal area relays semantic category information as well, or it is purely dictated by the shape of objects [7]–[10].

A parsimonious description of the emergence of category preferences from shape-selective responses would suggest that visual features distinguishing objects of different categories, presumably represented in IT cortex, can be used by classifiers in downstream regions such as prefrontal cortex to classify objects into semantic groups [7], [8], [11]. The prefrontal cortex (PFC) is also likely to play a critical role in integrating information received from upstream visual areas with memory to make task-relevant decisions and at the same time turn modulatory feedback signals to them [7], [12]. Consistent with this notion, multiple object recognition models have been suggested that successfully build complex representations of objects by hierarchically concatenating multiple layers of linear and non-linear operations, building a dictionary of shape features, which can then be decoded to categorical labels by a classifier [13]–[16]. Experiments controlling for the factor of shape have demonstrated that category-selective neural activity in IT is mostly due to shape, whereas prefrontal cortex shows malleable, task-dependent activity that can support categorization [7], [17]–[20].

Therefore, object recognition is likely to involve a wide range of sensory and higher order cognitive functions. While early visual areas are involved in the uni-modal processing of visual information, more advanced cognitive processes on visual information such as conscious perception, learning and decision making are performed at the level of higher cortical levels, such as the frontal cortex. While neural systems sub-serving these processes may be spatially far apart [3], [4], [18], [21], they require access to information from the early visual areas. How the brain areas responsible for these different levels of neural processing interact with one another during object recognition is poorly understood. Here, we focus in particular on how important the functional interaction between brain regions is for identification of object categories and how spatially dependent the role of these interactions in category identification is.

Previous studies have computed the strength of phase locking between activities of different brain regions, as proxy of neural communication between them [22]–[26]; however, it is not yet clear how functionally as well as behaviorally relevant this coupling may be for identification of object categories in human brain (see however [27] for a documentation of content-specific frontal-parietal phase coupling in monkeys). In order to examine these interactions, we recorded electrocorticography (ECoG) field potentials from epilepsy patients and measured the strength of phase locking between different brain areas [28]. We further investigated if this inter-areal synchrony is informative of different object categories presented to the human subjects. Our data show not only that the inter-areal synchrony pattern is category-dependent but that the spatial pattern of this category selectivity is behaviorally relevant, i.e., local connections with the occipital area and global connections with the frontal regions show the highest behavioral relevance of category-selectivity. These results highlight the anatomical dependence of the contribution of local neuronal circuitries vs. global connectivity patterns to the transmission of object category information in the human brain.

## Methods

### Experimental procedure

#### Participants

Participants were 10 people, aged between 12-34, diagnosed with pharmacologically intractable epilepsy. The recordings were performed in either Children’s Hospital Boston (CHB) or Brigham and Women’s Hospital (BWH), with the aim of localizing the seizure foci for potential surgical resection. All participants provided informed consent and were ensured about the confidentiality of their data. The study was approved by the institutional review boards at both hospitals. For more detailed information on the dataset, see the original publication on the data [28].

#### Recordings and Preprocessing

Intracranially implanted electrodes (Ad-Tech, Racine, WI, USA; each recording site was 2 mm in diameter with 1 cm separation) [29], [30] were utilized to localize the seizure event foci. Recording regions were set out in grids or strips, each containing 4 to 64 recording sites. The recorded sites ranged between 48 to 126 for each subject (80.4 ± 18.4, mean ± SD). All recorded neural signals were amplified (×2500), filtered between 0.1 and 100 Hz, and sampled with 256 Hz sampling rate at CHB (XLTEK, Oakville, ON, Canada) and 500 Hz at BWH (Bio-Logic, Knoxville, TN, USA). The data were further denoised using a 60 Hz notch filter and each electrode’s signal was globally referenced [28]. Although this referencing approach may suffer from global artifacts, it maintains neuronal activity relevant to the macroscopic (rather than mesoscopic) scale of brain state (see [31] for a mesoscopic approach). Subjects were hospitalized 6–9 days, during which physiological data were continuously monitored. The data presented in this study is based on the periods without seizure activity.

#### Stimulus Presentation

Objects from five different categories were presented as contrast-normalized grayscale images to subjects: ‘‘animals,’’ ‘‘chairs,’’ ‘‘human faces,’’ ‘‘fruits,’’ and ‘‘vehicles’’. Contrast normalization was done by fixing the grayscale pixel levels’ standard deviation. Images were shown for 200 ms separated by an interval of 600 ms, while subjects performed a one-back memory task to detect the repeated presentation of the same image category. A pseudorandom order was used to present the images.

### Data Analysis

#### Phase Locking Value (PLV)

Recorded data were filtered between 4-32 Hz in 4 Hz-wide bands with an overlap of 2 Hz using the EEGLAB toolbox [32] in MATLAB (The MathWorks, Natick, MA). For each frequency band, we calculated the PLV between the recorded signals from each pair of electrodes: A unit-length vector was assigned to each temporal sample with its phase equal to the phase difference between the two electrodes. These vectors were averaged over temporal samples, and finally the length of resultant vectors was averaged over trials (Equation 1):

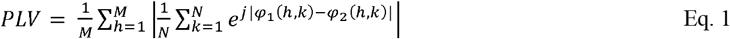

Where M is the number of trials, N is the number of temporal samples, and □_1_ and □_2_ are the instantaneous phase for the signals recorded from electrodes 1 & 2, derived using the Hilbert transform and |v| represents the length of vector v. PLV represents the magnitude of phase locking between the given electrode pair.

#### Category selectivity (RV)

To calculate the category selectivity of PLV, the across-category variation of PLV was divided by the average within-category variation of PLV (a measure called RV):

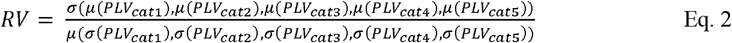

Where *σ* represents the standard deviation, *μ* denotes the average and cat_i_ stands for the *i*th category. Higher values of RV reflect a larger distinction between the different categories’ PLVs.

#### Behavioral relevance of category selectivity

To compute how different category selectivity is between hit and miss trials, for each category, RV was computed for a randomly selected subset of hit trials with the same size as the miss trials. This procedure was repeated for 50 iterations and the z-score of RVs for miss vs. hit trials was calculated:

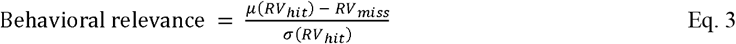

where RV_miss_ and RV_hit_ are computed over the set of miss trials and a given subset of hit trials, respectively and *μ*/*σ* are calculated over the randomly selected subsets of hit trials (with an equal size as the miss trials). This measure reflects how strongly category selectivity is linked to the subject’s behavioral performance.

The dependence of behavioral relevance on the distance between the areas was examined for each region by studying the lateral length of the connections from the lowest and highest octiles. To this end, a random selection of connections with the same size as the high and low behaviorally relevant connections were made for each area (for 100,000 repetitions). Next, the absolute difference between the highest and lowest behavioral relevance octiles was computed for each repetition and the original absolute difference was compared to that of the randomly generated data, generating a p-value by calculating the proportion of random differences that are less likely than the original difference value.

## Results

We examined whether the functional interaction between two different brain regions has any role in the encoding of distinct visual object categories. To investigate this, we recorded Electrocorticogram (ECoG) signals from different brain regions of epileptic patients, while they performed a one-back memory task involving different object categories. We calculated the degree of phase-locking between the intracranial field potentials (IFPs) of each pair of electrodes by measuring the phase locking value (PLV) between the recorded signals across different bands of the IFPs’ low frequency range (4-32Hz) (see Methods – Equation 1). Figure 1 shows example data from two different pairs of electrodes with high, and low PLVs (0.92 and 0.36, respectively). The location of each electrode pair is shown in the brain insets. The left panel representing a high PLV depicts how the phase difference between the two electrodes remains constant across time, whereas the two signals in the right panel do not maintain a fixed phase difference through time. This is an example of how variable the inter-areal functional interaction could be, motivating the question whether this variation contributes to the processing of object categories.

**Figure 1:**
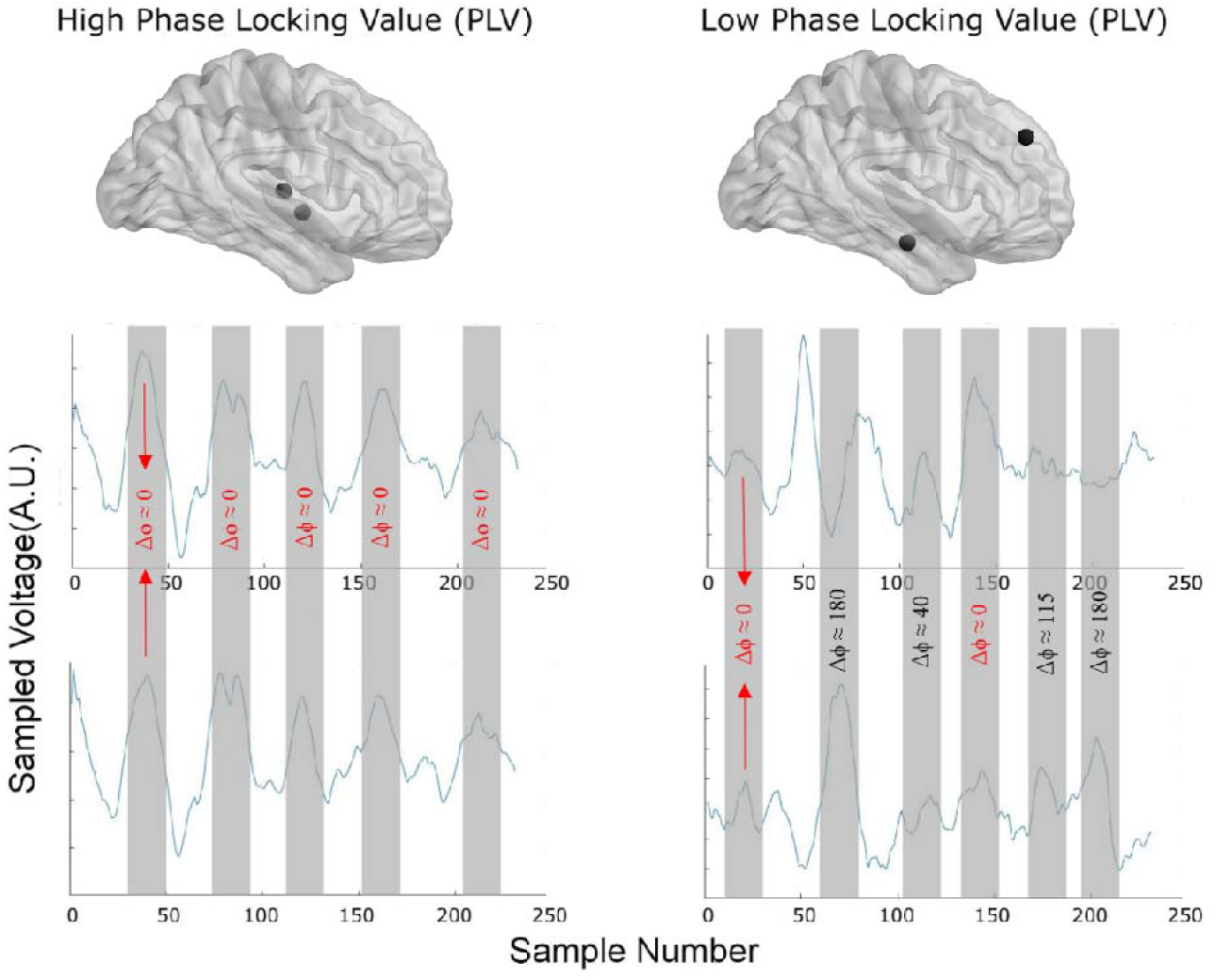
Example signals for different PLV magnitudes. Sample IFP signals recorded from two different pairs of electrodes with low (0.34) and high (0.90) Phase Locking Value (PLV). As highlighted, the peaks and troughs from the two recording electrodes are well aligned in the high PLV condition (left column), whereas no clear alignment is observed in the low phase locking condition (right column). Electrodes come from the middle temporal gyrus and superior lateral temporal gyrus (left) and medial Parahippocampal gyrus and middle frontal gyrus (right).

We next computed (for each frequency band) the category selectivity of each electrode pair’s PLV, by dividing the variation (across categories) of PLV by the mean of PLV variation within each category (see Methods - Equation 2). Figure S schematically depicts high and low category selectivity in two imaginary categories. To enable across-subject analysis of the data, we focused our analyses on only those regions which were recorded in at least two different subjects and averaged this category selectivity measure across the low frequency bands. We next computed the category selectivity for the PLV between all possible pairs of electrodes within each subject’s data. For each pair of brain regions then the category selectivity was averaged between the corresponding electrode pairs (Figure 2A). The strength of category selectivity for each pair of regions is color-coded, showing its variability within and across subjects. The analyzed regions (n=24) across the dataset are shown in the across-subject super map (Figure 2B). Observing the map of PLV-based category selectivity for individual subjects clearly visualizes how category selectivity varies across different pairs of brain regions.

**Figure 2:**
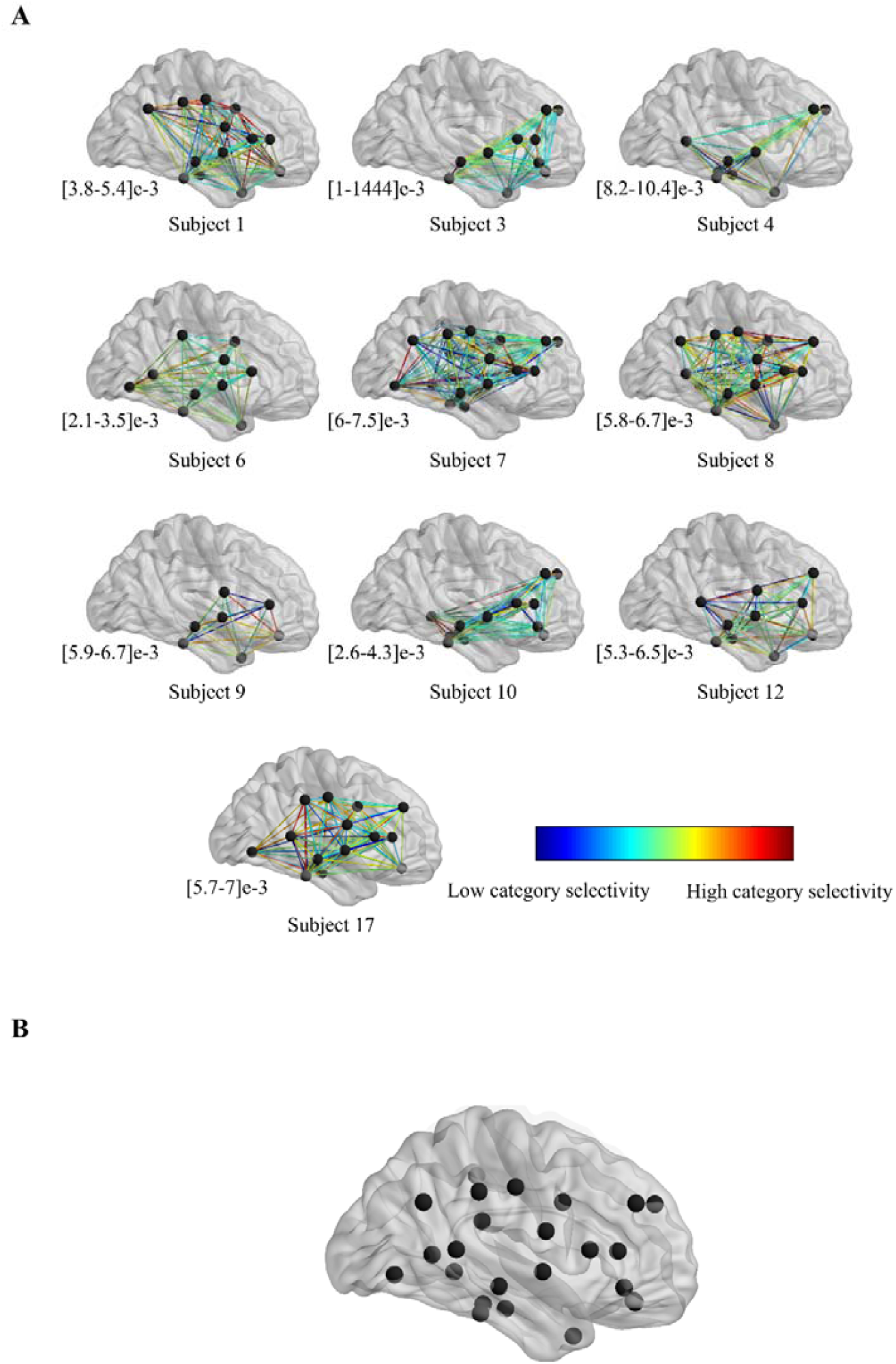
PLV-based category selectivity for different pairs of brain regions. **(A)** Brain maps show the location of regions at which at least one electrode was placed for each subject and the strength of PLV-based category selectivity for each pair of brain regions. **(B)** Location of analyzed brain regions (those regions that have been recorded in more than one subject [N=24]).

In the final step, we computed the behavioral relevance of RV, by randomly selecting a subset of hit trials equal to the number of wrongly performed (miss) trials in each category for 50 repetitions and calculating the z-score of RV for miss relative to hit trials (see Methods) (Figure 3A). Figure 3B shows the distribution of behavioral relevance across all possible edges of all subjects which is significantly above zero (p < 1e-78, sign test), indicating that category selectivity of inter-regional functional interaction needs to be large enough for the subjects to accomplish the trial successfully. These results suggest that not only the functional interaction between regions (reflected by PLV) is different across the presentation of different object categories, but also that this difference is relevant for the perceptual behavior of the subjects.

**Figure 3:**
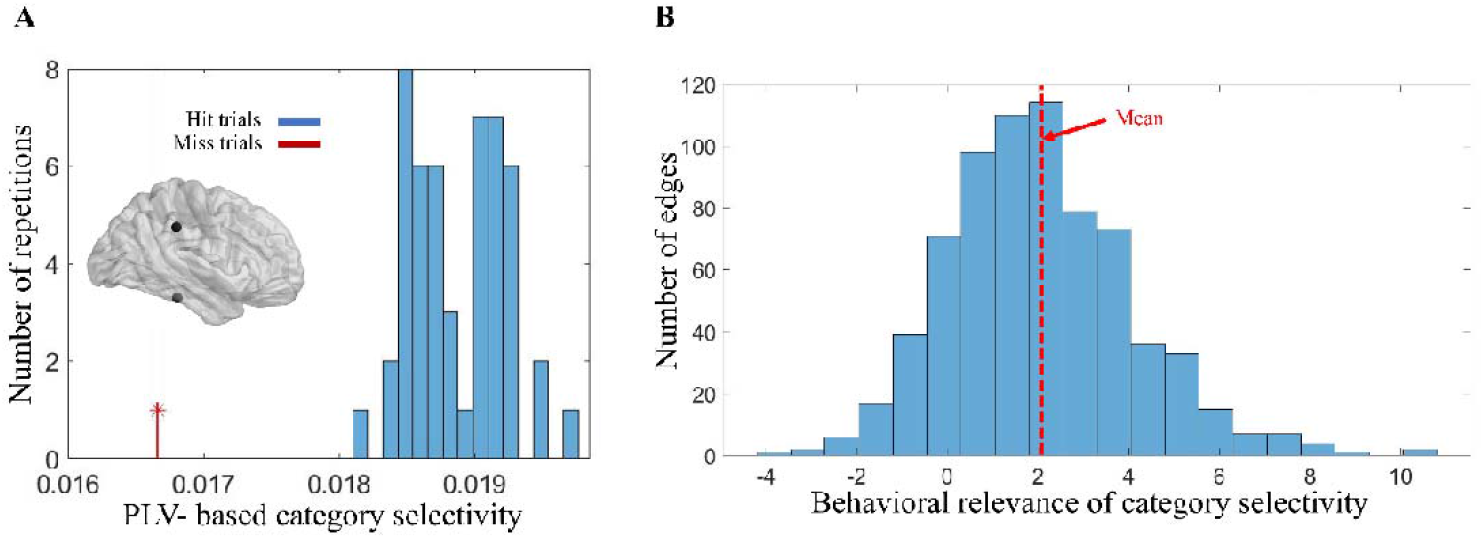
Distribution of category selectivity and its behavioral relevance for pairs of brain regions. **(A)** An example edge’s behavioral relevance (z-score = 6.4) for a sample subject; Each of the histogram’s data points represents the category selectivity (calculated using Eq. 2) for one randomly selected subset of hit trials (with the same number as the miss trials). The data point shown with a red line indicates the category selectivity computed for the miss trials (N = 309) of the sample subject (subject 6). Obviously, the miss trials’ category selectivity is considerably separated from that of the hit trials selected subsets. **(B)** Histogram of category selectivity’s behavioral relevance (Eq. 3) across significant and non-significant edges of all subjects with each data point representing the behavioral relevance for each of 715 edges (p< 1e-78, sign test).

To investigate the spatial organization of behaviorally relevant edges across the brain, we averaged each edge’s behavioral relevance across all subjects and consequently partitioned them into eight groups sorted by their behavioral relevance. We next selected the groups with the lowest and highest behavioral relevance. Figure 4A shows the two groups of edges categorized into the main anatomical brain lobes (“frontal,” “parietal,” “temporal,” “limbic” and occipital) based on either of their ending points. Figure 4B quantifies the contribution of each region to the high/low behavioral relevance network, by counting the number of edges from each region separated by behavioral relevance. For the frontal lobe it is obvious that the edges with the highest behavioral relevance are mostly those which connect different brain lobes extended across the brain volume (named global here), whereas the connections with the lowest behavioral relevance are more local (p-value = 0.03, permutation test). This effect is even more pronounced for electrodes in the prefrontal cortex (p-value = 0.009, permutation test). However, the opposite effect is observed for occipital (p-value = 0.006, permutation test) lobe (Figure 5). This is consistent with the proposal that the functional interaction of purely sensory areas (like occipital cortex) with closer neighboring areas is of more behaviorally crucial importance compared to the interaction with higher order areas. On the other hand, for a high-level associative area and particularly the prefrontal cortex (with a profound role in multi-modal processing), functional interaction (in terms of visual information) with the network of distantly located areas is of more behavioral relevance rather that interaction with anatomically close-by brain areas.

**Figure 4:**
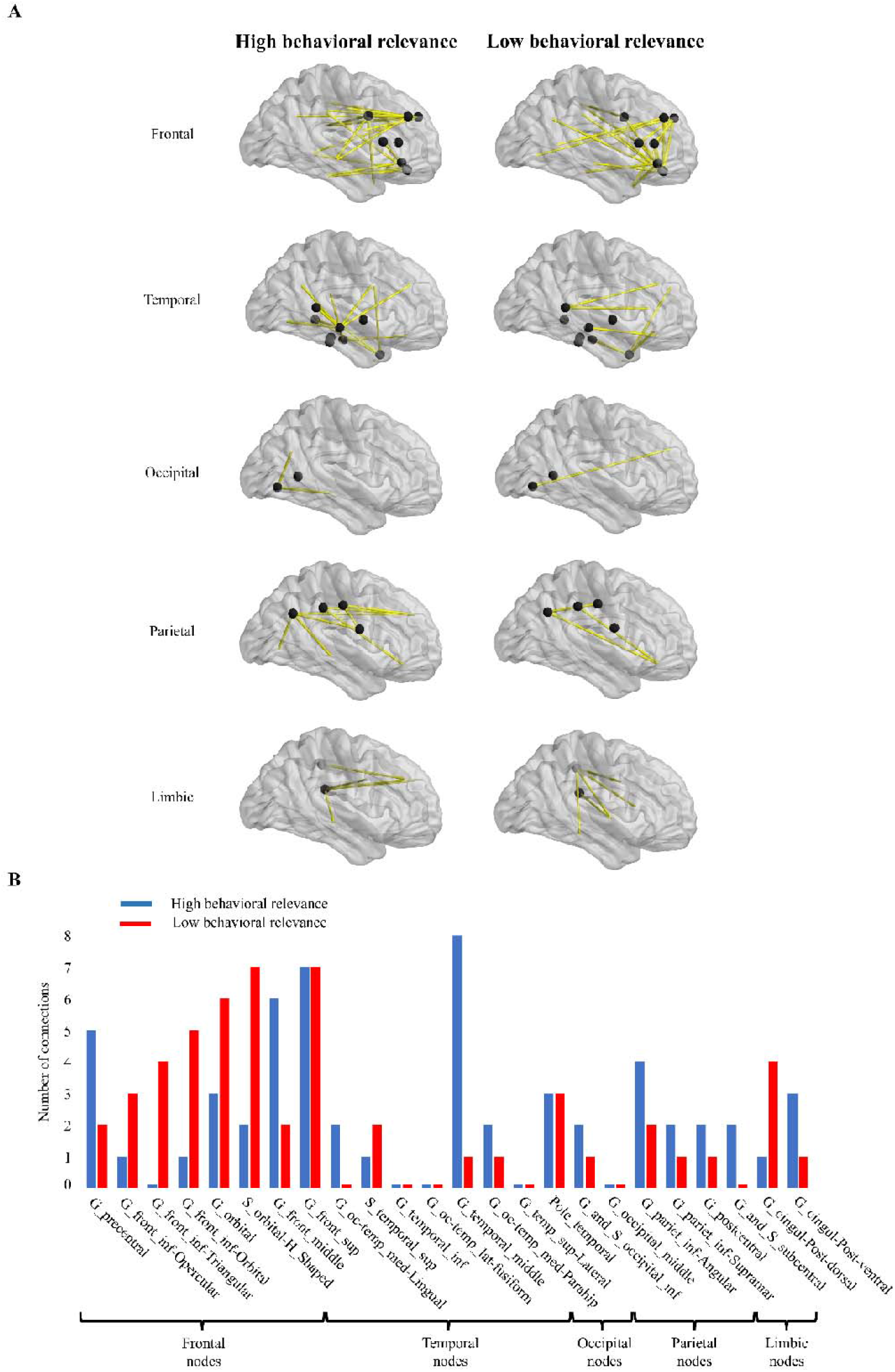
Different brain regions’ contribution to the behavioral dependence of connectivity. **(A)** Left and right columns represent connections from the highest and lowest quantile (octile), respectively of behavioral relevance, separated by involvement of each brain lobe. **(B)** Distribution of connections for each region, categorized by behavioral relevance, with high behavioral relevance nodes in blue and low behavioral relevance nodes in red.

**Figure 5:**
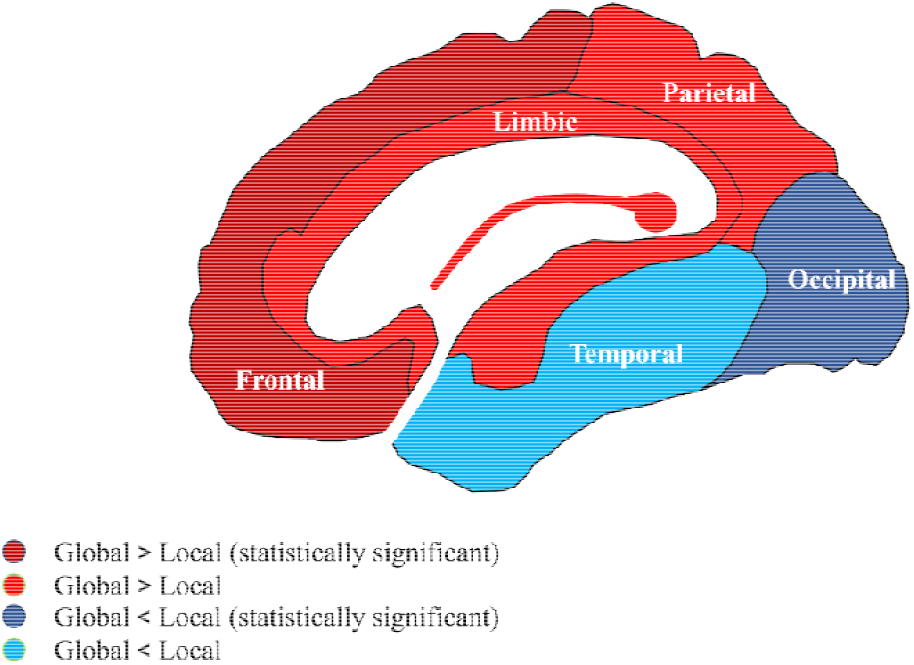
Behavioral relevance of category selectivity for each brain lobe. Lighter-colored regions are the regions with no significant p-values (parietal, temporal, and limbic), red indicates the regions with a stronger behavioral relevance for global compared to local connections, and blue in contrast shows a stronger behavioral relevance for local, compared to global connections.

## Discussion

The role of global inter-areal neuronal interactions vs. localized intra-areal processing in representation and conscious perception of object information has been widely emphasized but its behavioral relevance remains unclear [33]. Our data show that i) there is a phase locking between electrode (region) pairs, ii) the behavioral performances are linked to the category selectivity of the phase locking value, and iii) the information about the behavior is dependent on the length of the connections between the areas. Specifically, for the frontal lobe, the longer (global) connections are more responsible for the behavior of the subject; however, for the occipital lobe, the shorter (local) connections have the higher responsibility for producing the subjects’ behavior.

Normally, brain imaging studies on humans use imaging methods, such as fMRI, EEG, MEG, all suffering from a lack of either temporal resolution (as for fMRI) or spatial resolution (as for EEG and MEG). These limitations have been an obstacle for examining the fine time-resolved interaction of distant local brain regions. In this study, using ECoG electrodes, we recorded intracranially from a multitude of brain regions many reported to be involved in object categorization. This allowed us for the first time to address the behavioral relevance of inter-areal interactions for the representation of object categories. Our results are the first evidence for a potential role of inter-areal interaction in encoding of environmental object categories in the human brain, supporting the thesis that not only the single regions matter in object perception, but also their interaction plays an independent role in perceptual coding. While here we focused on the role of connections and the importance of region-dependent globality in object recognition, understanding the directionality of the interactions may help us to comment more precisely on the causal role of regional nodes especially in global connections.

Previous studies evaluated the role of information transfer and connections between different sensory and executive areas in the process of object recognition. In an experimental study Bar et al. (2006), tested a model that could describe the full path of recognizing the identity of an object in the brain using fMRI data. In this model after early analysis, information about low spatial frequency components of a picture is transferred in a fast pathway from primary visual areas in cerebral cortex to frontal cortex which is sent back to temporal cortex to be combined with other information for final decision [34].

Our finding of the distinguished behavioral role of frontal lobe’s connectivity with distantly located regions is also consistent with previous observations where an impairment of long frontal-temporal pathways severely affected object discrimination performance [35], [36]. Our result is further in-line with the prefrontal cortex’ unique setting, transmitting information to/from many distinct brain circuits and a critical role in a wide set of other critical cognitive abilities; attention, memory, decision making and perceptual binding [37]–[40]. Our observation aligns well with the existing evidence that emphasizes the importance of distributed neural networks for visual processing elucidating the distinct roles of frontal and occipital lobes in object recognition. The finding that global connections across brain lobes are more behaviorally relevant for the frontal lobe, while local connections are more crucial for the occipital lobe, is consistent with the hierarchical organization of the visual system.

Although we found that the behavioral relevance of local and global connections differs when addressing different brain regions, there is a consistent relationship between behavior and the length of connections, since we did not observe any significant length-dependence of behavioral relevance for connections with the limbic, temporal and parietal lobe.

In conclusion, our study has advanced our understanding of object recognition by illustrating the critical role of functional interactions between distinct brain regions. It was demonstrated that the category selectivity of these interactions, reflected by phase-locking values, not only varies across different object categories but also significantly influences the perceptual behavior of subjects. Moreover, we found that the behavioral relevance of these interactions is spatially organized in a way that underscores the unique roles of different brain areas in object recognition. Specifically, global connections extending across the brain volume are more behaviorally relevant for the frontal lobe, while local connections with closer neighboring areas are more crucial for the occipital lobe. These insights align with our understanding of the occipital cortex’s role in sensory processing and the prefrontal cortex’s role in multi-modal processing. Overall, these findings pave the way for more nuanced explorations of brain region interactions and their influence on object recognition and other cognitive functions.

## Supplementary Data

**Supplementary Figure 1:**
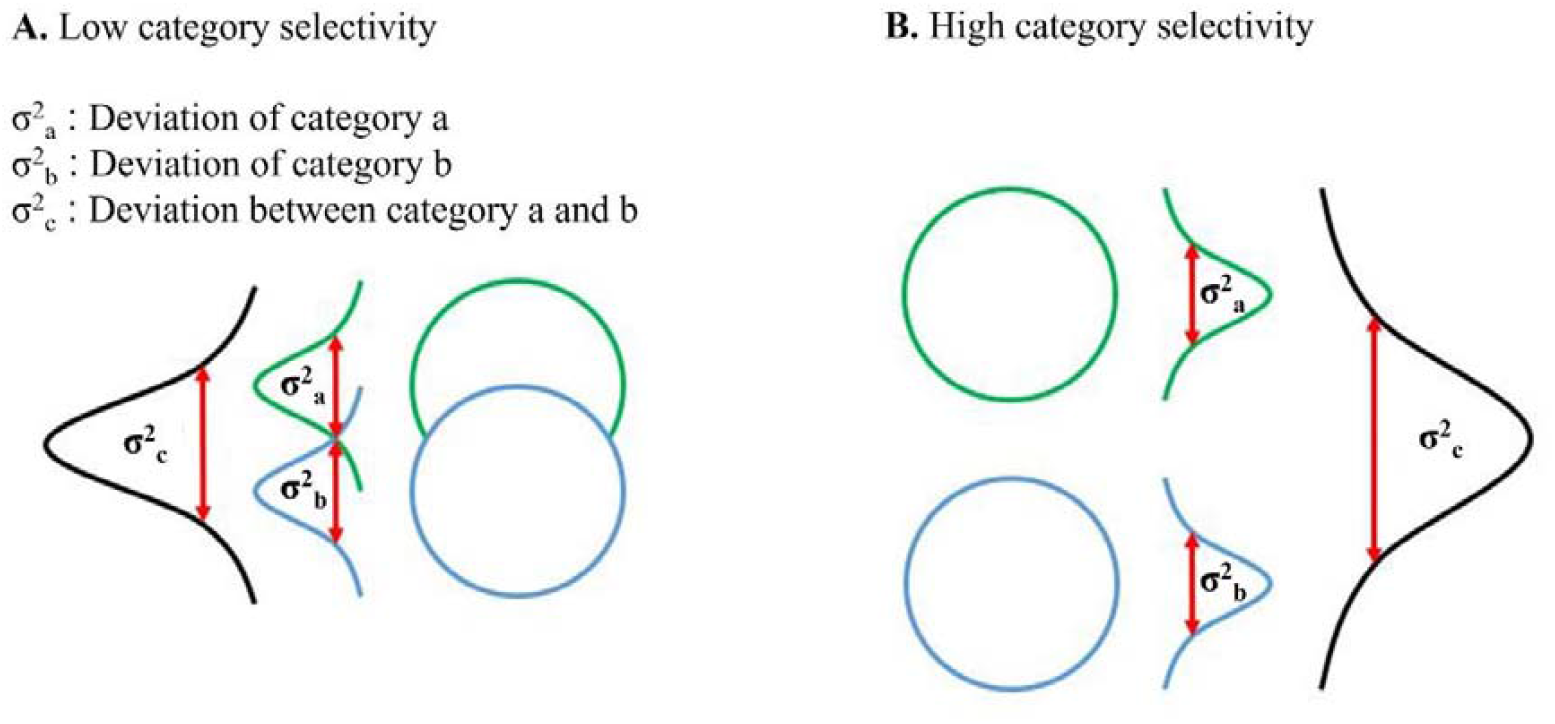
Schematic example of low and high category selectivity (measured using RV). A low RV value (left panel) indicates that the distributions corresponding to different categories are less distinguishable (larger intra-category variance relative to the across-category variance), while more distinct categories correspond to higher values of RV (right panel). Circles schematically represent the set of PLVs corresponding to each object category.

## References

[1] M. Cauchoix, S. M. Crouzet, D. Fize, and T. Serre, “Fast ventral stream neural activity enables rapid visual categorization,” NeuroImage, vol. 125, pp. 280–290, Jan. 2016, doi: 10.1016/j.neuroimage.2015.10.012.

[2] K. Grill-Spector and K. S. Weiner, “The functional architecture of the ventral temporal cortex and its role in categorization,” Nat. Rev. Neurosci., vol. 15, no. 8, Art. no. 8, Aug. 2014, doi: 10.1038/nrn3747.

[3] D. J. Freedman and J. A. Assad, “Neuronal Mechanisms of Visual Categorization: An Abstract View on Decision Making,” Annu. Rev. Neurosci., vol. 39, no. 1, pp. 129–147, 2016, doi: 10.1146/annurev-neuro-071714-033919.

[4] J. J. DiCarlo, D. Zoccolan, and N. C. Rust, “How does the brain solve visual object recognition?,” Neuron, vol. 73, no. 3, pp. 415–434, Feb. 2012, doi: 10.1016/j.neuron.2012.01.010.

[5] L. G. Ungerleider and A. H. Bell, “Uncovering the visual ‘alphabet’: Advances in our understanding of object perception,” Vision Res., vol. 51, no. 7, pp. 782–799, 2011, doi: 10.1016/j.visres.2010.10.002.

[6] A. H. Bell, F. Hadj-Bouziane, J. B. Frihauf, R. B. H. Tootell, and L. G. Ungerleider, “Object Representations in the Temporal Cortex of Monkeys and Humans as Revealed by Functional Magnetic Resonance Imaging,” J. Neurophysiol., vol. 101, no. 2, pp. 688–700, Feb. 2009, doi: 10.1152/jn.90657.2008.

[7] D. J. Freedman, M. Riesenhuber, T. Poggio, and E. K. Miller, “A Comparison of Primate Prefrontal and Inferior Temporal Cortices during Visual Categorization,” J. Neurosci., vol. 23, no. 12, pp. 5235–5246, Jun. 2003, doi: 10.1523/JNEUROSCI.23-12-05235.2003.

[8] C. Baldassi, A. Alemi-Neissi, M. Pagan, J. J. Dicarlo, R. Zecchina, and D. Zoccolan, “Shape similarity, better than semantic membership, accounts for the structure of visual object representations in a population of monkey inferotemporal neurons,” PLoS Comput. Biol., vol. 9, no. 8, p. e1003167, 2013, doi: 10.1371/journal.pcbi.1003167.

[9] N. Sigala, F. Gabbiani, and N. K. Logothetis, “Visual categorization and object representation in monkeys and humans,” J. Cogn. Neurosci., vol. 14, no. 2, pp. 187–198, Feb. 2002, doi: 10.1162/089892902317236830.

[10] A. Bardon, W. Xiao, C. R. Ponce, M. S. Livingstone, and G. Kreiman, “Face neurons encode nonsemantic features,” Proc. Natl. Acad. Sci., vol. 119, no. 16, p. e2118705119, Apr. 2022, doi: 10.1073/pnas.2118705119.

[11] S. Bracci and H. Op de Beeck, “Dissociations and Associations between Shape and Category Representations in the Two Visual Pathways,” J. Neurosci. Off. J. Soc. Neurosci., vol. 36, no. 2, pp. 432–444, Jan. 2016, doi: 10.1523/JNEUROSCI.2314-15.2016.

[12] E. K. Miller and J. D. Cohen, “An integrative theory of prefrontal cortex function,” Annu. Rev. Neurosci., vol. 24, pp. 167–202, 2001, doi: 10.1146/annurev.neuro.24.1.167.

[13] M. Riesenhuber and T. Poggio, “Hierarchical models of object recognition in cortex,” Nat. Neurosci., vol. 2, no. 11, Art. no. 11, Nov. 1999, doi: 10.1038/14819.

[14] T. Serre, A. Oliva, and T. Poggio, “A feedforward architecture accounts for rapid categorization,” Proc. Natl. Acad. Sci. U. S. A., vol. 104, no. 15, pp. 6424–6429, Apr. 2007, doi: 10.1073/pnas.0700622104.

[15] D. L. K. Yamins, H. Hong, C. F. Cadieu, E. A. Solomon, D. Seibert, and J. J. DiCarlo, “Performance-optimized hierarchical models predict neural responses in higher visual cortex,” Proc. Natl. Acad. Sci. U. S. A., vol. 111, no. 23, pp. 8619–8624, Jun. 2014, doi: 10.1073/pnas.1403112111.

[16] A. Krizhevsky, I. Sutskever, and G. E. Hinton, “ImageNet Classification with Deep Convolutional Neural Networks,” in Advances in Neural Information Processing Systems, Curran Associates, Inc., 2012. Accessed: Nov. 26, 2023. [Online]. Available: https://proceedings.neurips.cc/paper_files/paper/2012/hash/c399862d3b9d6b76c8436e924a68c45b-Abstract.html

[17] D. J. Freedman, M. Riesenhuber, T. Poggio, and E. K. Miller, “Visual categorization and the primate prefrontal cortex: neurophysiology and behavior,” J. Neurophysiol., vol. 88, no. 2, pp. 929–941, Aug. 2002, doi: 10.1152/jn.2002.88.2.929.

[18] D. J. Freedman, M. Riesenhuber, T. Poggio, and E. K. Miller, “Categorical representation of visual stimuli in the primate prefrontal cortex,” Science, vol. 291, no. 5502, pp. 312–316, Jan. 2001, doi: 10.1126/science.291.5502.312.

[19] J. L. McKee, M. Riesenhuber, E. K. Miller, and D. J. Freedman, “Task dependence of visual and category representations in prefrontal and inferior temporal cortices,” J. Neurosci. Off. J. Soc. Neurosci., vol. 34, no. 48, pp. 16065–16075, Nov. 2014, doi: 10.1523/JNEUROSCI.1660-14.2014.

[20] E. M. Meyers, D. J. Freedman, G. Kreiman, E. K. Miller, and T. Poggio, “Dynamic Population Coding of Category Information in Inferior Temporal and Prefrontal Cortex,” J. Neurophysiol., vol. 100, no. 3, pp. 1407–1419, Sep. 2008, doi: 10.1152/jn.90248.2008.

[21] E. K. Miller, D. J. Freedman, and J. D. Wallis, “The prefrontal cortex: categories, concepts and cognition.,” Philos. Trans. R. Soc. B Biol. Sci., vol. 357, no. 1424, pp. 1123–1136, Aug. 2002, doi: 10.1098/rstb.2002.1099.

[22] T. Womelsdorf et al., “Modulation of neuronal interactions through neuronal synchronization,” Science, vol. 316, no. 5831, pp. 1609–1612, Jun. 2007, doi: 10.1126/science.1139597.

[23] C. A. Bosman et al., “Attentional stimulus selection through selective synchronization between monkey visual areas,” Neuron, vol. 75, no. 5, pp. 875–888, Sep. 2012, doi: 10.1016/j.neuron.2012.06.037.

[24] P. Fries, “A mechanism for cognitive dynamics: neuronal communication through neuronal coherence,” Trends Cogn. Sci., vol. 9, no. 10, pp. 474–480, Oct. 2005, doi: 10.1016/j.tics.2005.08.011.

[25] X. Wang et al., “Representing object categories by connections: Evidence from a mutivariate connectivity pattern classification approach,” Hum. Brain Mapp., vol. 37, no. 10, pp. 3685–3697, May 2016, doi: 10.1002/hbm.23268.

[26] S. Jafakesh, F. Z. Jahromy, and M. R. Daliri, “Decoding of object categories from brain signals using cross frequency coupling methods,” Biomed. Signal Process. Control, vol. 27, pp. 60–67, May 2016, doi: 10.1016/j.bspc.2016.01.013.

[27] R. F. Salazar, N. M. Dotson, S. L. Bressler, and C. M. Gray, “Content-specific fronto-parietal synchronization during visual working memory,” Science, vol. 338, no. 6110, pp. 1097–1100, Nov. 2012, doi: 10.1126/science.1224000.

[28] H. Liu, Y. Agam, J. R. Madsen, and G. Kreiman, “Timing, timing, timing: fast decoding of object information from intracranial field potentials in human visual cortex,” Neuron, vol. 62, no. 2, pp. 281–290, Apr. 2009, doi: 10.1016/j.neuron.2009.02.025.

[29] A. K. Engel, C. K. E. Moll, I. Fried, and G. A. Ojemann, “Invasive recordings from the human brain: clinical insights and beyond,” Nat. Rev. Neurosci., vol. 6, no. 1, pp. 35–47, Jan. 2005, doi: 10.1038/nrn1585.

[30] M. J. Kahana, R. Sekuler, J. B. Caplan, M. Kirschen, and J. R. Madsen, “Human theta oscillations exhibit task dependence during virtual maze navigation,” Nature, vol. 399, no. 6738, Art. no. 6738, Jun. 1999, doi: 10.1038/21645.

[31] J. Wang, A. Tao, W. S. Anderson, J. R. Madsen, and G. Kreiman, “Mesoscopic physiological interactions in the human brain reveal small-world properties,” Cell Rep., vol. 36, no. 8, p. 109585, Aug. 2021, doi: 10.1016/j.celrep.2021.109585.

[32] A. Delorme and S. Makeig, “EEGLAB: an open source toolbox for analysis of single-trial EEG dynamics including independent component analysis,” J. Neurosci. Methods, vol. 134, no. 1, pp. 9–21, Mar. 2004, doi: 10.1016/j.jneumeth.2003.10.009.

[33] G. A. Mashour, P. Roelfsema, J.-P. Changeux, and S. Dehaene, “Conscious Processing and the Global Neuronal Workspace Hypothesis,” Neuron, vol. 105, no. 5, pp. 776–798, Mar. 2020, doi: 10.1016/j.neuron.2020.01.026.

[34] M. Bar et al., “Top-down facilitation of visual recognition,” Proc. Natl. Acad. Sci., vol. 103, no. 2, pp. 449–454, Jan. 2006, doi: 10.1073/pnas.0507062103.

[35] P. G. F. Browning, A. Easton, and D. Gaffan, “Frontal-temporal disconnection abolishes object discrimination learning set in macaque monkeys,” Cereb. Cortex N. Y. N 1991, vol. 17, no. 4, pp. 859–864, Apr. 2007, doi: 10.1093/cercor/bhk039.

[36] D. Gaffan, A. Easton, and A. Parker, “Interaction of inferior temporal cortex with frontal cortex and basal forebrain: double dissociation in strategy implementation and associative learning,” J. Neurosci. Off. J. Soc. Neurosci., vol. 22, no. 16, pp. 7288–7296, Aug. 2002, doi: 10.1523/JNEUROSCI.22-16-07288.2002.

[37] P. Andrés, “Frontal cortex as the central executive of working memory: time to revise our view,” Cortex J. Devoted Study Nerv. Syst. Behav., vol. 39, no. 4–5, pp. 871–895, 2003, doi: 10.1016/s0010-9452(08)70868-2.

[38] S. M. Courtney, L. Petit, J. M. Maisog, L. G. Ungerleider, and J. V. Haxby, “An area specialized for spatial working memory in human frontal cortex,” Science, vol. 279, no. 5355, pp. 1347–1351, Feb. 1998, doi: 10.1126/science.279.5355.1347.

[39] S. M. Szczepanski, C. S. Konen, and S. Kastner, “Mechanisms of Spatial Attention Control in Frontal and Parietal Cortex,” J. Neurosci., vol. 30, no. 1, pp. 148–160, Jan. 2010, doi: 10.1523/JNEUROSCI.3862-09.2010.

[40] M. Parto Dezfouli, P. Schwedhelm, M. Wibral, S. Treue, M. R. Daliri, and M. Esghaei, “A neural correlate of visual feature binding in primate lateral prefrontal cortex,” NeuroImage, vol. 229, p. 117757, Apr. 2021, doi: 10.1016/j.neuroimage.2021.117757.

